# Leaf metabolic traits reveal hidden dimensions of plant form and function

**DOI:** 10.1101/2023.03.08.531692

**Authors:** Tom W. N. Walker, Franziska Schrodt, Pierre-Marie Allard, Emmanuel Defossez, Vincent E. J. Jassey, Meredith C. Schuman, Jake M. Alexander, Oliver Baines, Virginie Baldy, Richard D. Bardgett, Pol Capdevila, Phyllis D. Coley, Nicole M. van Dam, Bruno David, Patrice Descombes, Maria-Jose Endara, Catherine Fernandez, Dale Forrister, Albert Gargallo-Garriga, Gaёtan Gauser, Sue Marr, Steffen Neumann, Loïc Pellissier, Kristian Peters, Sergio Rasmann, Ute Roessner, Roberto Salguero-Gómez, Jordi Sardans, Wolfram Weckwerth, Jean-Luc Wolfender, Josep Peñuelas

## Abstract

The plant metabolome encompasses the biochemical mechanisms through which evolutionary and ecological processes shape plant form and function^1,2^. However, while the metabolome should thus be an important component of plant life-history variation^3^, we know little about how it varies across the plant kingdom. Here, we use the plant functional trait concept^4^ – a powerful framework for describing plant form and function^5–7^ – to interpret leaf metabolome variation among 457 tropical and 339 temperate plant species. Distilling metabolite chemistry into five discriminant metabolic functional traits reveals that plants vary along two major axes of leaf metabolic specialization – a leaf chemical defense spectrum and an expression of leaf longevity. These axes are qualitatively consistent for tropical and temperate species, with many trait combinations being viable. However, axes of leaf metabolic specialization vary orthogonally to life-history strategies described by widely used functional traits^5–7^, while being at least equally important to them. Our findings question classical trait^6^ and plant defense^8^ theory that predicts relationships between the leaf chemical phenotype, plant productivity, and pace of life. Moreover, we show that metabolic functional traits describe unique dimensions of plant life-history variation that are complementary to, and independent from, those captured by existing plant functional traits.

## Main Text

Plants produce a staggering diversity of metabolites – upwards of one million throughout the plant kingdom and many thousands within an individual^9^ (the metabolome). The plant metabolome is well known to human society, being the source of most cosmetics^10^, flavors^11^, and medicines^12^. Yet, little is known about how the metabolome varies across the plant kingdom. The metabolome is the biochemical basis of physiology, comprising primary metabolites involved in cellular function (e.g., carbohydrates, amino acids) and specialized metabolites produced to confront stress (e.g., flavonoids, terpenoids)^9,13,14^. The metabolome thus encompasses the biochemical mechanisms through which evolutionary and ecological processes shape plant form and function (i.e., morphology and physiology ^7^)^1,2^. Plant form and function vary predictably among species because evolutionary and ecological processes generally act to maximize fitness^8,15–18^, thus limiting the number of morphological or physiological strategies that are viable^5–7^. It follows that the metabolome should also be constrained to a limited number of strategies that maximize fitness^3^. However, although cross-species studies are now emerging^19–22^, no attempt has been made to characterize how the metabolome varies systematically throughout the plant kingdom. We thus remain unable to place the vast diversity of the plant metabolome into a coherent ecological context.

The plant functional trait concept offers a powerful framework to contextualize the plant metabolome. Plant functional traits are a standardized set of morphological and physiological characteristics that provide universal metrics of plant form and function^4^. Plant functional traits have been pivotal in identifying the life-history strategies that govern plant fitness^5–7^, understanding plant impacts on population^23^, community^24^, and ecosystem^25^ processes, and predicting global change effects on ecosystems^26^. Plant functional traits thus provide a strong reference point from which to interpret variation in the plant metabolome. However, measurements of the metabolome remain absent from the functional trait concept, in part because they yield complex data describing thousands of metabolites from hundreds of biochemical pathways^3^. Advances in chemoinformatics make it now possible to translate metabolite identities into a targeted set of chemical properties^27,28^. Doing so resonates with the functional trait concept – i.e., that quantitative characteristics provide a bridge between identity and function^4^ – and is an intuitive way to contextualize the metabolome against plant functional traits. Yet, while chemoinformatics is commonplace in natural products chemistry^29–31^, it remains entirely absent from ecology.

Contextualizing the plant metabolome against existing plant functional traits raises two alternative hypotheses. If the metabolome varies colinearly with functional traits, then it offers a biochemical validation of the functional trait concept. Such validation is necessary because functional traits are emergent properties of multiple biochemical processes^4,32^, making them proxies for physiology that can be poor predictors of ecological processes^33–35^. Alternatively, if the metabolome varies orthogonally to functional traits, then it describes dimensions of plant life-history variation missed by existing functional traits. Such orthogonal variation would enhance the explanatory power of the functional trait concept by providing additional axes of trait specialization to examine plant form and function. Either way, integrating measurements of the plant metabolome into the plant functional trait concept will yield insight into how the metabolome varies across the plant kingdom, while also offering a high-resolution mechanistic lens with which to advance the functional trait concept itself.

Here, we performed an assessment of the leaf metabolomes of 457 tropical and 339 temperate plant species to characterize variation in the metabolome and test whether it validates, or expands upon, existing plant functional traits. First, we examined the chemical properties of all identified metabolites (N = 4292) to derive a minimum set of “metabolic functional traits” that capture chemical variation in the leaf metabolome. We then characterized how metabolic traits vary among species to identify major axes of leaf metabolic specialization in the plant kingdom. Finally, we combined metabolic traits with eight widely used functional traits (plant height, seed mass, stem specific density, leaf area, specific leaf area and leaf carbon, nitrogen and phosphorus concentrations; hereafter “classical traits”)^7^ to determine whether axes of metabolic specialization are colinear with, or orthogonal to, plant life-history strategies described by classical functional traits.

### Five chemical properties capture chemical variation in the leaf metabolome

We expressed the leaf metabolomes of tropical^36^ and temperate^37^ plant species as the presence or absence of annotated metabolic features (i.e., metabolites) detected through liquid chromatography mass spectrometry (tropical N = 3356; temperate N = 2227). We then characterized the chemistry of the leaf metabolome by examining variation in the chemical properties of all unique metabolites from both datasets (N = 4292)^31^. We focused on 21 properties describing metabolite constitution, geometry, or topology that have relationships with (bio)chemical activity (Table S1). Pairwise correlations among chemical properties (Fig. 1A, upper panel) reveal that properties separate into five clusters, each representing a distinct facet of leaf metabolite chemistry. The largest cluster (Fig. 1A, green) describes overall size and structural complexity, comprising total numbers of atoms and bonds, molecular weight, a predictor of polarity based on the proportion of non-carbon atoms (M log *P*)^38^, and related topological parameters (i.e., Wiener numbers^38^, Eccentric Connectivity^39^). A second cluster (Fig. 1A, purple) describes bond conjugation and reactivity, comprising numbers of aromatic atoms and bonds, numbers of atoms in the largest conjugated (pi-) system, and a derived complexity index indicative of promiscuous activity in drugs (*γ*_MF_)^40^. A third cluster (Fig. 1A, yellow) describes polar intermolecular (non-covalent) forces, comprising total and mass-specific polar surface area and numbers of hydrogen (H) bond donors or acceptors^41,42^. Such forces are important for the selective binding of small molecules to DNA^43^, RNA^44^, and proteins^45^. A fourth cluster (Fig. 1A, blue) comprises two predictors of metabolite polarity based on summed properties of atoms and bonds (X log *P*^46^, A log *P*^47^), where lower values indicate higher polarity. A final cluster (Fig. 1A, red) describes carbon bond saturation *via* the relative occurrence of sp^3^ (unsaturated, three-dimensional) and sp^2^ (saturated, planar) hybridization^48^. Overall, these clusters indicate that leaf metabolite chemistry is highly structured and can be described by five distinct facets: size/complexity, bond conjugation/reactivity, contributions to polar intermolecular forces, general polarity, and carbon bond saturation.

**Figure 1.**
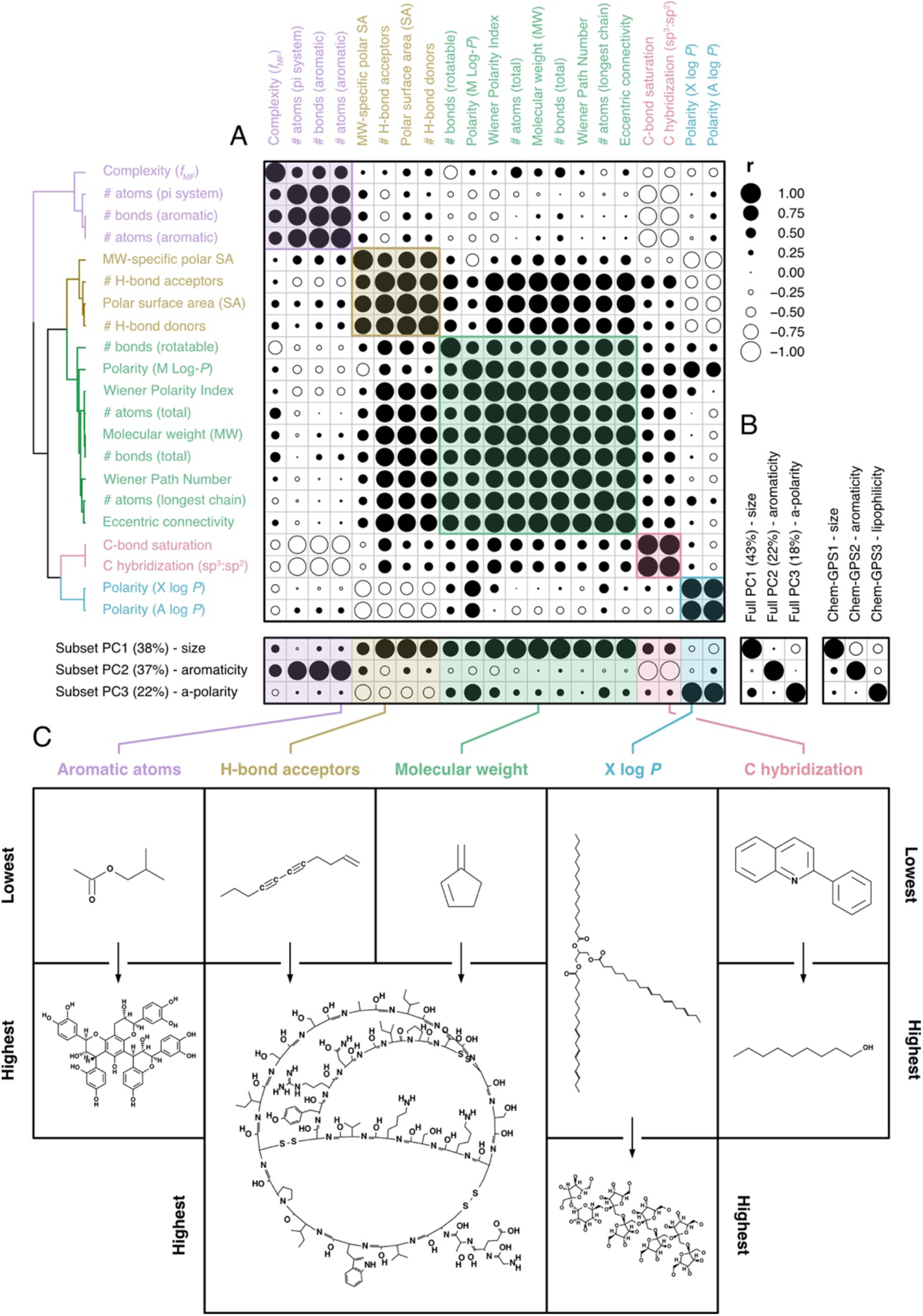
Variation in leaf metabolite chemistry can be captured by five chemical properties. (A) Coefficients for Pearson correlations among 21 chemical properties (Table S1) of annotated metabolites detected in tropical and temperate datasets (N =4292). Properties are ordered based on correlation similarity (hierarchical clustering; Ward), with colors separating distinct clusters of properties (maximal silhouette width: K = 5). The lower panel shows correlations between chemical properties and scores from the first three axes of a PCA performed on a subset of five properties, as well as (B) between subset PC scores and either scores from a PCA performed on all 21 properties (left panel) or metabolite positions on the first three axes of ChemGPS-NP^31^ space (right panel). Circle sizes indicate coefficient strengths, while fills discriminate between positive (black) and negative (white) correlations. (C) The chemical properties selected to represent each cluster, alongside chemical graphs of metabolites with the lowest and highest values for each trait.

The strong collinearity among chemical properties (Fig. 1A, upper panel) suggests that the five facets of leaf chemistry can be represented by a small number of discriminant variables. Indeed, a PCA performed on a subset of five chemical properties, one from each cluster (Fig. 1C), is equivalent to a PCA performed on all 21 properties (Fig. 1B, left panel; PC1: r_4290_ = 0.97; PC2: r_4290_ = 0.99; PC3: r_4290_ = 0.93; *P* < 0.001 in all cases). We selected molecular weight, H-bond acceptor count, aromatic atom count, polarity (X log *P*), and carbon hybridization (sp^3^:sp^2^) as five properties that are intuitive for non-specialists to interpret (Fig. 1C; Table S1). However, most combinations of properties covering all facets of leaf chemistry also yield equivalent PCAs (Fig. S4). Pairwise correlations between all properties and subset PC scores further confirm that selected properties capture observed leaf chemical variation, in that correlations are consistent within each cluster (Fig. 1A, lower panel).

Selected properties also describe biologically relevant chemical space according to an existing framework that classifies natural products based on size, then aromaticity, then lipophilicity (Fig. 1B, right panel; PC1: r_4290_ = 0.99; PC2: r_4290_ = 0.87; r**4290** = 0.81; *P* < 0.001 in all cases)^31^. The first axis of the subset PCA (38% variance) separates heavy metabolites with many H-bond acceptors from light metabolites with fewer H-bond acceptors. Alignment between metabolite size and numbers of H-bond acceptors is expected because larger molecules usually engage in stronger intermolecular interactions^49^. The second axis (37% variance) separates reactive metabolites with more aromatic carbon bonds from unreactive metabolites with more saturated carbon bonds. This axis is also intuitive because conjugated bond systems are unsaturated, reactive, and planar, whereas saturated bonds, such as those abundant in fatty acids, are unreactive and three-dimensional^50^. The third axis (18% variance) separates nonpolar metabolites from polar metabolites^29^. Our findings thus show that leaf chemical variation can be represented by five chemical properties, one from each facet of leaf chemistry.

### Five chemical properties provide more information than metabolite family

The five selected chemical properties not only represent leaf metabolite chemistry (Fig. 1), but also reflect known differences among metabolite families. Using the subset PCA to interpret variation among metabolite families (Fig. 2), we observe that peptides are large, display average bond saturation, and are moderately polar; carbohydrates are of average size, display high bond saturation, and are highly polar; and fatty acids are smaller, display high bond saturation, and are of average polarity. Lignans, alkaloids, coumarins and flavonoids, families of specialized metabolites, display low bond saturation due to the presence of many aromatic bonds, with alkaloids and coumarins also being relatively small. Terpenoids, a family of abundant specialized metabolites (N = 2306), vary in size, bond saturation, and polarity, echoing their diversity across the plant kingdom^51^. Although chemical properties generally differentiate among metabolite families, all families possess variable and overlapping chemistries (Fig. 2, marginal boxplots). We thus suggest that chemical properties, quantitative metrics of the chemistry that underpins metabolite composition and activity, provide more precise information about metabolite function than metabolite family identity. Indeed, drug discovery and medicinal chemistry rely on many of the same chemical properties to screen for bioactive molecules^29–31^. Such a perspective resonates with the functional trait concept by reaching beyond taxonomy to describe function^4^ and provides a strong foundation for a novel set of metabolic functional traits.

**Figure 2.**
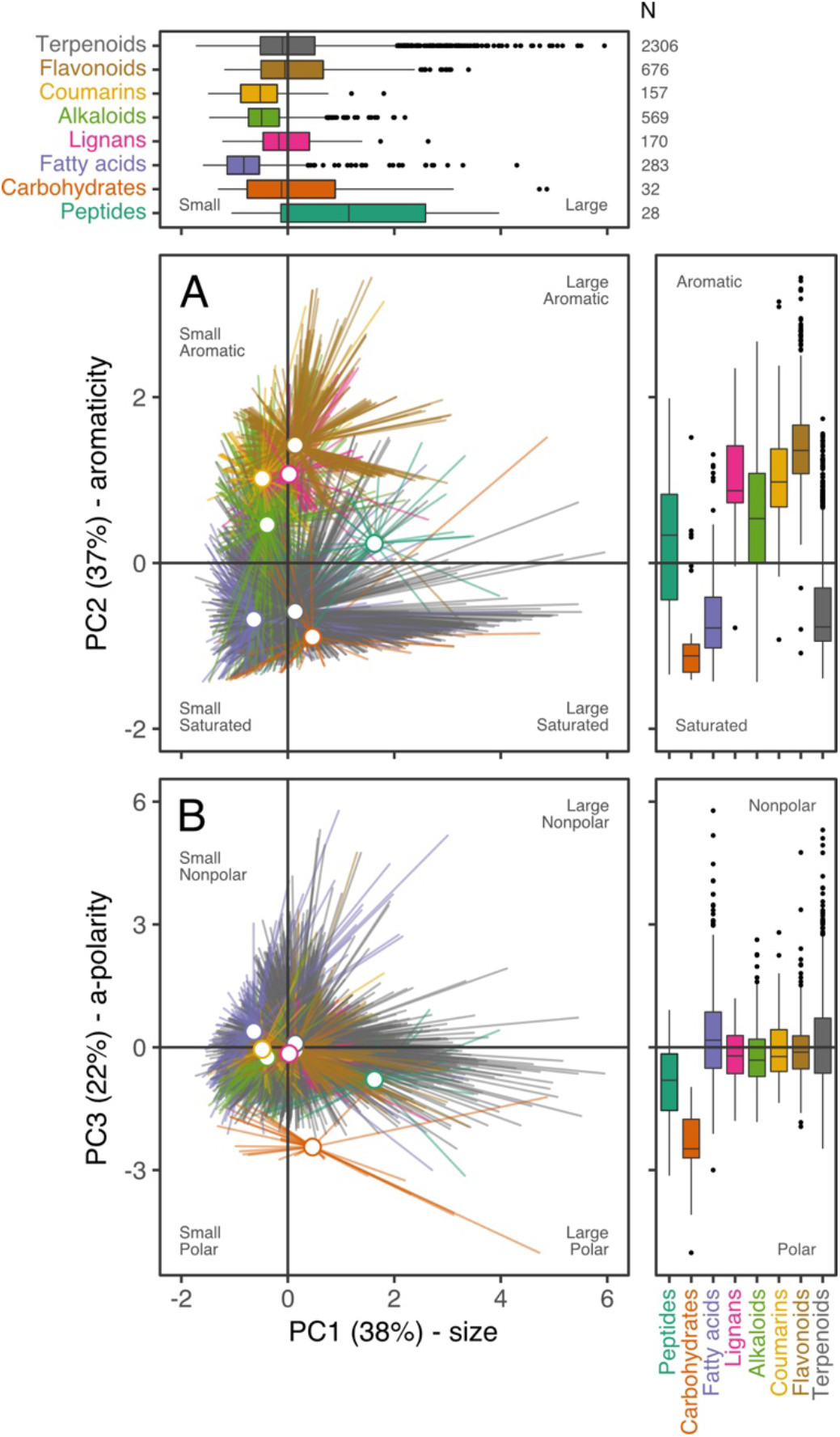
Five chemical properties describe variation among and within metabolite families. Biplots of PC1 scores and (A) PC2 scores or (B) PC3 scores from a PCA describing variation in selected chemical properties (C hybridization, H-bond acceptors, molecular weight, polarity, aromatic atoms) among all annotated leaf metabolites (N = 4221) detected in 862 plant species. The PCA is functionally equivalent to a PCA containing all 21 chemical properties (see Main Text), with axes separating leaf metabolites based on size (PC1), aromaticity (PC2), and a-polarity (PC3; see Fig. 1A, lower panel). Colors separate metabolite families as defined by marginal boxplot fills/labels (center: median; box: 25% to 75% quantiles; whiskers: 1.5 × IQR; points: outliers beyond whiskers).

### Five metabolic functional traits describe two axes of leaf metabolic specialisation

We converted selected properties into plant-level metabolic functional traits to examine how the leaf metabolome varies among tropical and temperate plant species. We first compared the volume and “lumpiness” (i.e., aggregation around values) of multidimensional space occupied by metabolic traits to four null models to test whether species converge on a restricted set of interrelated trait combinations^7^. Observed hypervolumes of metabolic traits are 98.1% to 99.8% smaller and 71.0% to 94.7% lumpier than those of three null models assuming no trait covariance (Table S2; Table S3; always *P* < 0.001 in all cases). However, observed hypervolumes are larger (tropical: *P* = 0.002; temperate: *P* = 0.047) than that of the null model assuming trait covariance and selection against extreme values, while also being similarly lumpy (tropical: *P* = 0.662; temperate: *P* = 0.858). By comparison, observed hypervolumes of eight classical traits are smaller than null models assuming no trait covariance (Table S2; *P* < 0.001 in all cases), either the same size as (tropical species; *P* = 0.154) or smaller than (temperate species; *P* = 0.018) that of the null model assuming selection against extreme values, and lumpier than all null models (Table S3; *P* < 0.001 in all cases). Collectively, these findings yield three insights. First, metabolic functional traits are interrelated, so plants are constrained to a limited number of viable metabolic trait combinations. Second, while extreme metabolic trait values may be selected against in temperate species, this is not so for tropical species. Expressing extreme leaf metabolome chemistry is thus a viable life-history strategy in tropical environments, potentially due to more intense biotic interactions selecting for a diversity of defensive leaf metabolites and thus divergence of chemical strategies^52^. Third, metabolic traits are less aggregated around non-extreme values than classical traits. As such, although plants converge towards one of several viable classical trait strategies (e.g., woody, non-woody^7^), they can possess a wider array of leaf metabolome strategies.

We performed PCAs on metabolic traits to characterize major axes of leaf metabolic specialization among species. Metabolic trait variation is consistent among tropical and temperate species, with 92% variation being explained by the first two PCs (Fig. 3). The first axis (tropical: 52%; temperate 49%) separates species that produce more unsaturated, aromatic, nonpolar leaf metabolites from those that produce more saturated, polar, leaf metabolites. This axis likely describes a spectrum of leaf chemical defense strategies^8^. Unsaturated aromatic metabolites, such as alkaloids, coumarins, and flavonoids (Fig. 2), are reactive and serve as toxins or antioxidants in response to stress. For instance, conjugated bond structures can interfere with protein function by binding covalently to sidechains^53^, generate or quench oxidative stress^54^, or absorb damaging wavelengths of light^55^. By contrast, saturated non-aromatic metabolites can play different roles in leaf defense, both directly as toxins^56^ and indirectly as signaling molecules^57^. An alternative explanation is that the first axis reflects an investment (or not) into defense metabolites^8^, which could arise due to the high cost of synthesizing aromatic metabolites^17^. However, all plants must defend their leaves against stress, and we find that plants can maximize fitness at either end of this axis (Fig. 3). It is thus more plausible that the first axis represents a leaf chemical defense spectrum.

**Figure 3.**
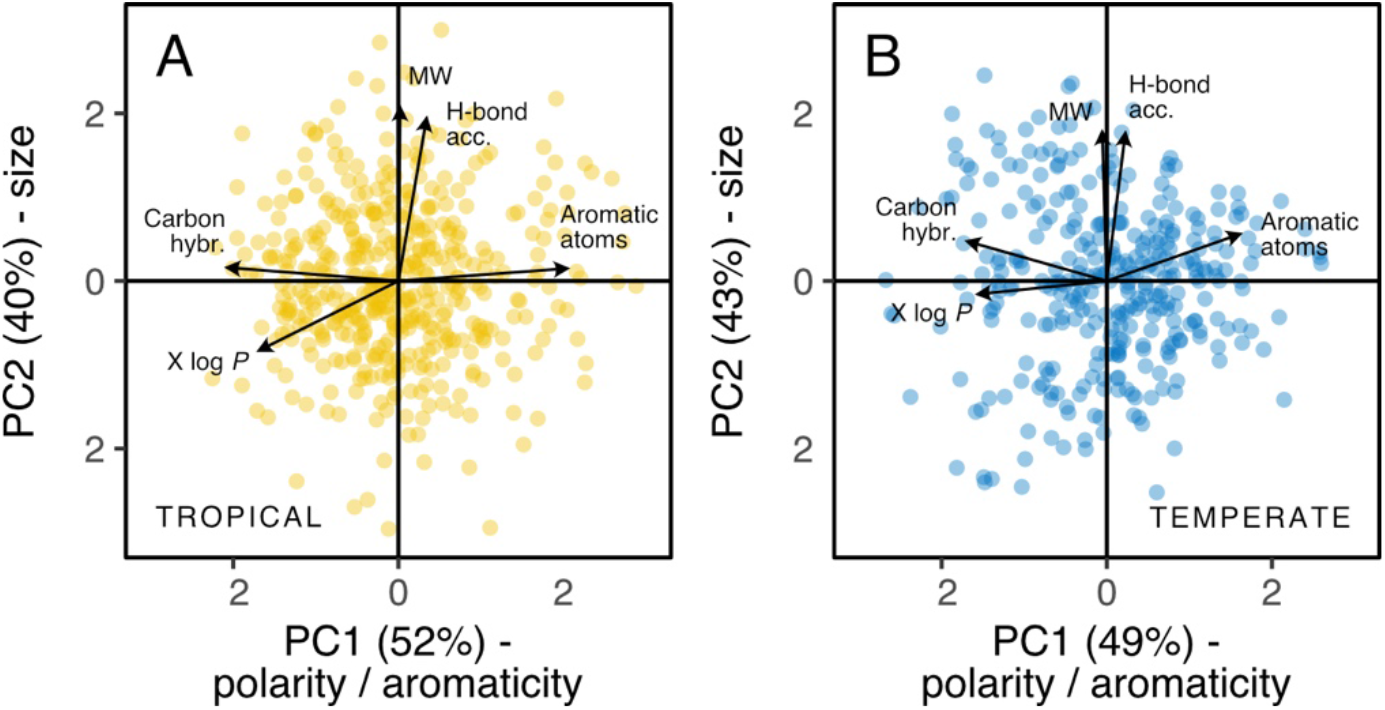
Metabolic traits describe two axes of leaf metabolic specialization among plant species. Biplots of PC1 and PC2 scores for PCAs describing variation in five metabolic functional traits (arrows) between (A) tropical (yellow; N = 457) and (B) temperate (blue; N = 339) plant species. Axes separate species based on the mean aromaticity *versus* polarity (inverse X log *P*) and carbon bond saturation (PC1) and mean size/complexity (PC2) of all annotated leaf metabolites present (Methods).

The second PCA axis (tropical: 40%; temperate: 43%) discriminates species that produce larger metabolites with many H-bond acceptors from those that produce smaller metabolites with fewer H-bond acceptors. This axis likely relates to leaf longevity, since large metabolites, such as lignans, peptides, and waxes, form the basis of long-lived physical^58,59^ and storage^60^ structures. Many of these structures also depend on strong intermolecular forces to form stable macromolecules or complexes^61,62^, which are enhanced by H-bond donors and acceptors^42^. As for the first axis, plants can maximize fitness at either end of the second axis, with species at the positive end of the axis investing in metabolites indicative of a longer leaf lifespan. As such, the second axis is probably a manifestation of the fast-slow continuum^63^ at the leaf level. Alternatively, the second axis may reflect an investment (or not) into structurally complex metabolites for physical defense, such as lignans or waxes^58,59^. However, such metabolites act by creating rigid structures that lengthen leaf lifespan anyway^64^, so physical protection is one component of leaf longevity. Taken together, metabolic trait PCAs reveal two major axes of leaf metabolic specialization. These axes relate to metabolite bond saturation and polarity (i.e., a leaf chemical defense spectrum) and metabolite size and propensity for intermolecular interactions (i.e., leaf longevity), with many combinations yielding successful life-history strategies.

### Axes of leaf metabolic specialisation reveal hidden dimensions of plant form and function

The apparent consistency of metabolic trait variation with classical life-history theory^5–8,17,63,64^ raises the possibility that metabolic traits describe aspects of plant form and function already captured by classical traits. We thus performed a PCA on metabolic traits plus the same widely used functional traits used above to test whether axes of leaf metabolic specialization are colinear with, or orthogonal to, trait-based life-history strategies. Considered alone, classical trait variation supports life-history theory^7^, with the first two PCs (tropical: 49%; temperate: 51%) describing the leaf economics spectrum (SLA, leaf nitrogen, leaf phosphorus)^6^ and variation indicative of plant longevity (plant height, seed mass)^63^ and hardiness (stem density, leaf carbon)^5^ (Extended Data Fig. 1). However, when combining classical and metabolic traits in the same PCA, the first four PCs (tropical: 66%; temperate: 67%) are equivalent to axes described by individual PCAs (Fig. 4A, lower marginal matrices). The first and third axes describe the first two axes of leaf metabolic specialization (Fig. 3), whereas the second and fourth axes describe the first two axes of classical trait specialization (Extended Data Fig. 1). Pairwise correlations among all traits further show that metabolic and classical traits do not covary (Fig. 4A,B), but separate into six clusters describing unique aspects of either metabolic or classical trait variation. As such, major axes of leaf metabolic specialization vary orthogonally to major axes of classical trait variation (Fig. S4) and are at least equally relevant for describing variation in plant form and function among tropical and temperate plant species.

**Figure 4.**
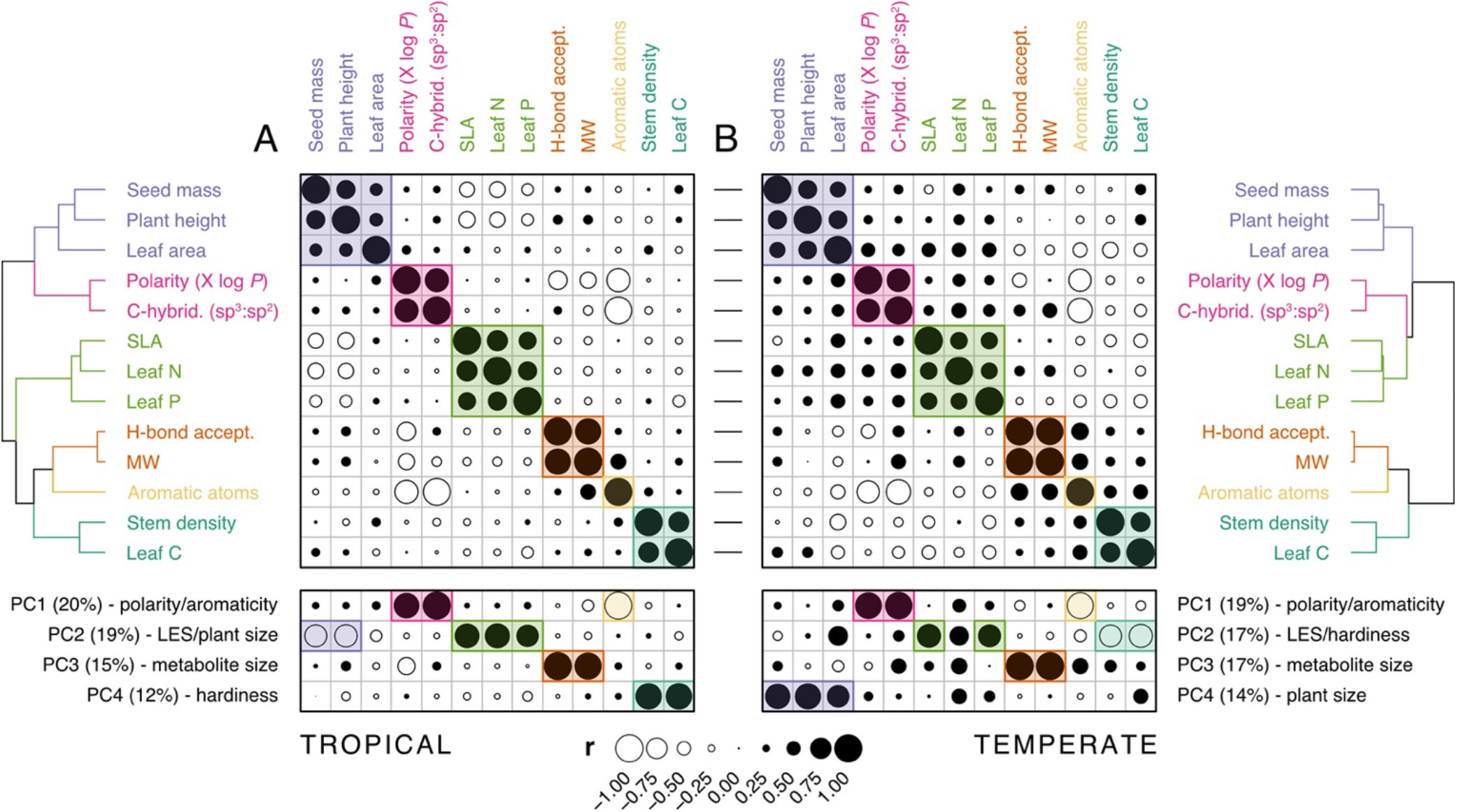
Metabolic and classical traits describe unique dimensions of plant form and function. Correlation coefficients for pairwise Pearson correlations among metabolic (Fig. 3) and classical functional traits (Extended Data Fig. 1) in (A) tropical (N = 457) and (B) temperate (N = 339) species. Traits are ordered based on correlation similarity (hierarchical clustering; Ward), with colors separating distinct clusters (maximal silhouette width: K = 6) and horizontal lines between matrices showing correspondence between tropical and temperate traits. Lower panel shows correlations between traits and scores from the first four axes of a PCA of the same traits (tropical: 66%; temperate: 67%; LES = leaf economics spectrum). Circle sizes indicate coefficient strengths, while fills discriminate between positive (black) and negative (white) correlations.

Leaf metabolism underpins energy production *via* photosynthesis^65^, but in doing so yields tissue that is sensitive to light^66^ and temperature^67^ and is of high nutritional value to herbivores^64^. Plants are thus under selective pressure to maximize photosynthetic rates while also protecting leaf tissue against damage. While this trade-off is consistent with life-history trade-offs derived from classical traits^5–7^, it does not necessarily follow that individuals apply the same strategy in every organ to maintain fitness. Indeed, we observe that species with “productive” trait values (e.g., high SLA, high leaf nitrogen) can produce leaves containing many complex aromatic metabolites, species possessing “hardy” trait values (e.g., high stem density, high leaf carbon) can produce leaves containing many simple polar metabolites, and species possessing trait values typical of a fast pace of life (e.g., low stature, small seeds) can produce leaves with many large stable or structural metabolites. Our findings thus question classical trait^6^ and plant defense^8^ theory that predicts relationships between the leaf chemical phenotype, plant productivity, and pace of life. In short, metabolic functional traits describe unique dimensions of plant life-history variation that are complementary to, and independent from, those captured by classical functional traits.

### Concluding remarks

In summary, we show that leaf metabolite chemistry is highly structured and can be described by five chemical properties. Using chemical properties as metabolic functional traits reveals that plants vary on two axes of leaf metabolic specialization – a leaf chemical defense spectrum (i.e., bond saturation, polarity) and leaf longevity (i.e., size, propensity for intermolecular interactions). Axes of metabolic specialization are similar in tropical and temperate species, with many combinations of trait values yielding successful life-history strategies. Axes are also consistent with expectations from life-history theory that plants invest strategically in tissue defense^6,8,17^ and between a fast *versus* slow pace of life^7,63^. Nevertheless, we find that metabolic trait variation is orthogonal to, not colinear with, classical trait variation. Metabolic functional traits thus reveal dimensions of plant life-history variation missed by existing functional traits. It is important to note that metabolomics data were acquired differently for tropical and temperate species and were matched to open functional trait databases. Nevertheless, we find consistent variation in the leaf metabolomes of 457 tropical and 339 temperate species that possess distinct geographic distributions and evolutionary histories (Extended Data Fig. 1A,B), as well as that encompass most documented functional trait variation (Extended Data Fig. 1G, red ribbons). We thus propose that distilling the leaf metabolome into metabolic functional traits captures macroecological patterns of metabolic variation widely across the plant kingdom, enhances the explanatory power of the functional trait concept, and offers a new set of tools for the discovery of species or genotypes with trait combinations adapted to societal needs.

## Methods

### Sample collection & preparation

We used two existing datasets describing the leaf metabolomes of tropical and temperate plant species (as the unit of observation). The tropical dataset originated from a random subsample of tropical leaves from the Pierre Fabre Sample Library^68^, which is an archive of ~17000 plant samples from *in situ* communities around the globe that was originally collected for drug discovery and is a registered collection of the European Union (registration code: 03-FR-2020)^69^. Samples were oven-dried (55 °C for 3 d) and extracted using overnight maceration (8 g sample; 80 mL ethyl acetate). Extracts were dried once (Genevac, SP Industries, Warminster, PA, USA) and eluted (30 mg extract plus 200 mg silica; 1g Silica SPE cartridges; 6 mL DCM plus 85:15 DCM:MeOH), then dried again and resuspended in DMSO at a final concentration of 5 mg mL^-1^. The temperate dataset originated from a study of landscape-scale ecological variation in phytochemical diversity^37^, wherein leaves were sampled from *in situ* grassland communities across Switzerland. Briefly, leaves were dried (40 °C for 5 d), milled (Retsch TissueLyser; Qiagen, Hilden, Germany) and extracted (20 mg tissue; 0.5 mL 80:19.5:0.5 MeOH:H2O: H2CO2). Extracts were homogenized (glass beads; 30 Hz for 3 min) and centrifuged (14000 RPM for 3 min), and the supernatant was isolated for measurement.

### Metabolomics measurements

Untargeted metabolomics was performed on all samples using Ultra High-Performance Liquid Chromatography Mass Spectrometry (UHPLC-MS). Tropical extracts were analyzed using a Waters Acquity UHPLC system coupled to a Q-Exactive Focus MS (Thermo Scientific, Bremen, Germany) with a heated electrospray ionization (HESI-II) source in positive mode. Extracts (2 μL) were separated on an Acquity C18 column (50 mm × 2.1 mm, 1.7 μm; Waters, Milford, MA, USA) at a flow rate of 600 μl min^-1^ using a binary solvent system (A: H2O plus 0.1% H_2_CO_2_; B: acetonitrile plus 0.1% H_2_CO_2_) and a linear gradient elution (5%-100% B, 7 min) plus isocratic elution (100% B, 1 min). MS data were acquired in data-dependent acquisition mode, in which MS/MS scans were performed on the three most intense ions detected during repeated full MS scans (full scans = 35000 FWHM at m/z 200; MS/MS scans = 17000 FWHM; MS/MS isolation window width = 1 Da; normalized collision energy = 15, 30 and 45 units). The MS was calibrated using a quality control (QC) mix containing caffeine, methionine-arginine-phenylalanine-alanine-acetate (MRFA), sodium dodecyl sulphate, sodium taurocholate and Ultramark 1621 dissolved in acetonitrile/methanol/H_2_O plus 1% H_2_CO_2_. Temperate extracts were analyzed using a Waters Acquity UHPLC system coupled to a Synapt G2 MS (Waters, Milford, MA, USA) with a heated electrospray ionization (HESI-II) source in positive mode. Extracts (2.5 μL) were separated on an Acquity C18 column (50 mm × 2.1 mm, 1.7 μm; Waters, Milford, MA, USA) at a flow rate of 600 μl min^-1^ using a binary solvent system (A: H_2_O plus 0.05% H_2_CO_2_; B: acetonitrile plus 0.05% H_2_CO_2_) and a linear gradient elution (2%-100% B, 6 min). MS data were acquired in data-independent acquisition mode, in which all precursor ions across the full mass range (85-1200 Da) were fragmented to yield MS/MS spectra. The MS was calibrated using a QC mix derived from all sample extracts.

### Metabolomics data pre-processing

Tropical MS data were treated using MZMine (version 2.53)^70^. Peak detection was performed using the centroid mass detector (MS noise = 1 × 10^4^; MS/MS noise = 0) and the ADAP chromatogram builder (min. scan group size = 5, min. group intensity = 1 × 10^4^; min. highest intensity = 5 × 10^5^; m/z tolerance = 12 ppm)^71^. Peaks were deconvoluted using the wavelets algorithm in ADAP with an intensity window SN (S/N threshold = 10; min. feature height = 5 × 10^5^; coefficient/area threshold = 130; peak duration range = 0-5 min; retention time (RT) wavelet range = 0.01-0.03 min). Isotopes were detected using the isotopes peak grouper (m/z tolerance = 12 ppm; absolute RT tolerance = 0.01 min absolute; max. charge = 2) and, where present, the most intense isotope was chosen. Peaks were filtered to retain features possessing an MS/MS scan and a RT of between 0.5 and 8 min, following which they were aligned using the join aligner method (m/z tolerance = 40 ppm; absolute RT tolerance = 0.2 min; m/z weight = 2; RT weight = 1). Temperate MS data were treated using MS-DIAL^72^. Peaks were detected with a minimum height of 300 and data were collected with an MS1 tolerance of 0.05 Da and an MS2 tolerance of 0.01 Da from 100–1200 Da and 0.5 to 12 minutes. Peaks were deconvoluted with a sigma window value of 0.5 and an MS/MS abundance cut-off of 50. Finally, raw features were aligned using the QC mix with a retention time tolerance of 0.1 minute. Following this, we separately clustered tropical and temperate metabolic features into sets of spectrally similar “consensus” features using molecular networks constructed on the Global Natural Products Social (GNPS; precursor and MS/MS fragment ion mass tolerances = 0.02 Da)^73,74^. We retained edges with a cosine score of more than 0.7 and with more than six matched peaks, permitted edges that connected two nodes only if each node appeared in the neighbour’s “top 10” most similar node list and, where necessary, limited the total size of a cluster to the 100 highest scoring edges. Finally, we converted peak height/area data to presence-absence data by summing peak heights/areas at the consensus metabolic feature level and expressing them as a one (present) or zero (absent) (tropical N = 7343, temperate N = 6682).

### Metabolite annotation

We annotated all metabolic features with chemical information by *in silico* spectral matching and taxonomically-informed scoring using a theoretically fragmented version of the Dictionary of Natural Products^75^ following^76,77^, retaining matches with a minimum score of 0.7 and at least six matched peaks. We assigned consensus metabolic features with a single consensus chemical classification, which we derived by taking the most common chemical classification across individual features contributing to a consensus feature. We filtered data to retain those features that were annotated with a metabolite identifier (i.e., SMILES notation; tropical N = 3356 (46%), temperate N = 2227 (33%)). We then derived the chemistry of annotated metabolites by using SMILES identifiers to query the Chemistry Development Kit^27^ with the package “rcdk”^78^. The CDK comprises 51 chemical descriptors, but we focused on 21 chemical properties that describe metabolite constitution, geometry, and topology (Table S1). We also derived values for annotated metabolites from the first three dimensions of the ChemGPS-NP tool^28^, which is an existing framework for classifying natural products that separates molecules based on size, then aromaticity, then lipophilicity^31^.

### Classical functional traits

All further data processing, analysis and visualization were undertaken in R^79^. For core functionality, we used the R packages “cowplot”^80^, “data.table”^81^, “drake”^82^, “furrr”^83^, “future”^84^, “parallel”^79^, “psych”^85^ and “tidyverse”^86^ (see below for packages pertaining to specific steps). Plant species names were cleaned against the Catalogue of Life and Encyclopedia of Life using the global name resolver in the package “taxize”^87^, following which we added a taxonomic hierarchy using “taxonlookup”^88^. We removed 14 tropical (ferns, cycads, conifers) and nine temperate (ferns, trees) species that were phylogenetically distinct from others within their respective datasets to prevent a small number of divergent species from overwhelming downstream analyses (Fig. S1). We obtained data for classical functional traits with proven relevance to plant form and function from the open-access version of the TRY database (version 5)^89^ – namely plant height (m), seed mass (mg), stem specific density (mg mm^-3^), leaf area (mm^2^), specific leaf area (mm^2^ mg^-1^) and leaf carbon, nitrogen and phosphorus concentrations (mg g^-1^)^6,7,90^. After removing clear outliers (reported error risk ≥ 4; values falling outside of documented limits), we calculated species-wise means and gap-filled missing trait values for all species present in the dataset (N = 46905 species) using a two-step process. First, we performed a Bayesian hierarchical probabilistic matrix factorization imputation (package “BHPMF”^91^), which is a taxonomically-constrained gap-filling approach that has been regularly applied to the TRY database^7,24^. The imputation was repeated 90 times, each time starting with a different combination of parameters (per-fold samples = 900-1000; cross-validation steps = 10-20; burn-in steps = 10% data length) and filtering out extreme (>1.5 times the maximum observed value for a trait) or highly uncertain (coefficient of variation >1) gap-filled values. We then calculated means of remaining gap-filled values across all iterations and used these to replace missing cases in original trait data. Second, we performed five iterations of multivariate imputation by chained equations (package “mice”^92^) on the partially gap-filled dataset and replaced remaining missing cases with the mean values from all iterations. Following this, we filtered fully gap-filled datasets for selected tropical and temperate species (see Table S4 for details about missing data), using genus-level mean trait values where species-level data were not available (215 tropical species, 5 temperate species). It is important to note that trait data coverage differed substantially between tropical and temperate species (Table S4). While this undoubtedly reduced the strength of observed trait variation in tropical relative to temperate data, it is unlikely that the reduced resolution of the tropical data changed the nature of functional trait variation, since artificially reducing the resolution of trait datasets to genus level did not change resulting PC axes (Fig. S2). Finally, we accounted for non-normal distributions in plant height, seed mass and leaf area data using log_10_-transformations and summarized trait variation among tropical and temperate species separately using scores from the first two components of varimax-rotated PCAs performed on all traits (Extended Data Fig. 1).

### Species distributions

We matched species names to Global Biodiversity Information Facility (GBIF) IDs using “rgbif”^93^ and gathered all available species occurrences from the GBIF website (www.gbif.org; tropical N = 336632, temperate N = 1847095). Working with each species individually, we filtered occurrences to include human observations and living or preserved specimens recorded from the year 1945 to present day and accompanied by either no individual counts or counts with a reasonable value (>100)^94^. We used the packages “CoordinateCleaner”^95^ to discard records falling in the sea, outside the stated country or within 10 km of capital cities, GBIF headquarters or known biodiversity institutions, records lying more than five times the interquartile range of the minimum distances to the nearest neighbor and records containing known issues with longitude/latitude information (i.e., zeros, rounding errors or conversion errors from other coordinate systems)^94,95^. Following this, we manually removed a further 20 tropical and four temperate records representing single occurrences within a continent, leaving a total of 239223 and 1251500 cleaned occurrences for tropical and temperate species, respectively. We plotted cleaned occurrences to visualize the geographical extents of tropical and temperate plant species (Extended Data Fig. 1AB), although it is important to emphasize that samples for metabolomics analyses were taken from within natural (not ornamental) distributions.

### Leaf metabolite chemistry

We collated all unique annotated metabolites detected in any tropical or temperate species into a single dataset capturing overall variation in the chemical properties of the leaf metabolome (N = 4292). We tested for interrelatedness between metabolite chemical properties using pairwise Pearson correlations coupled with hierarchical clustering (Ward; Fig. 1, upper panel), and estimated the number of discrete clusters of chemical properties using the maximal average silhouette width^96^ (Fig. 1A, dendrogram). We described major dimensions of leaf chemical variation using a varimax-rotated PCA performed on a subset of five representative chemical properties (one from each cluster: C-hybridization ratio, H-bond acceptor count, molecular weight, polarity (X log *P*), aromatic atom count). The first three PC axes described 97% variation, which we anchored back to individual chemical properties using pairwise Pearson correlations (Fig. 1A, lower panel). We tested whether the subset PCA was functionally equivalent to a full PCA performed on all 21 properties using two approaches. First, we performed pairwise Pearson correlations between subset *versus* full PC scores to determine whether the two PCAs separated metabolic features similarly on the same dimensions of chemical variation (Fig. 1B, left panel; see Main Text for statistics). Second, we used a Procrustes Rotation test (package “vegan”^97^; 999 permutations) to determine whether maximally-rotated configurations of full *versus* subset PCAs were significantly (i.e., non-randomly) associated (*m_12_* = 0.04, r^2^ = 0.98, *P* < 0.001). We also confirmed that the similarity among subset and full PCAs was not sensitive to which chemical properties were selected from each cluster. This was achieved by repeating Procrustes Rotation tests and PC score correlations for PCAs performed on every combination of five chemical properties that covered all five clusters and plotting distributions of resulting coefficients (Fig. S4). We performed pairwise Pearson correlations between subset PC scores and values from the first three dimensions of the ChemGPS-NP framework^28^ to test for correspondence between the subset PCA and existing descriptors of biologically relevant chemical space (Fig. 1B, right panel; see Main Text for statistics). We visualized chemical variation among metabolites using biplots of PC scores from the subset PCA, grouped by biochemical class (Fig. 2). We also illustrated the range of chemical structures captured by the five selected chemical properties by using the PubChem sketcher (https://pubchem.ncbi.nlm.nih.gov//edit3/index.html) to predict chemical graphs of metabolites with the lowest and highest values for each property (Fig. 1C).

### Plant-level metabolic functional traits

We selected C-hybridization ratio, H-bond acceptor count, molecular weight, X log *P*, and aromatic atom count to convert into five plant-level metabolic functional traits. For each species (i.e., unit of observation), we calculated the mean value of a chemical property considering all the metabolites present. In doing so, we generated datasets akin those capturing classical functional trait variation among tropical or temperate species, but instead quantifying variation in the mean C-hybridization ratio, H-bond acceptor count, molecular weight, polarity, and aromatic atom count of the annotated leaf metabolome among tropical or temperate species. We considered tropical and temperate datasets as separate entities for data analysis of trait variation, for four reasons. First, while *in silico* data treatment was identical for aligned MS data, sample collection and LC-MS measurements were not (see above), making it challenging to combine data on species-level metabolic variation without inflating differences between them. Second, unlike genomics and proteomics analyses, where only one type of molecule is isolated, untargeted metabolomics aims to simultaneously isolate thousands of molecules with a wide range of chemical properties^3,98^. Technical decisions made during sample extraction and analysis discriminate for or against certain types of metabolites, which filters the resulting view of the metabolome. There is still no single accepted method for extracting samples or acquiring metabolomics data^99^, so we chose here to use two datasets generated using different approaches to confirm that findings are not sensitive to the methods chosen. Third, classical functional trait and species occurrence data were sparser for tropical than temperate species (see *Classical Functional Traits,* above), so we chose to keep datasets separate rather than artificially reduce the resolution of the temperate dataset. Fourth, maintaining a separation between the two datasets increased our interpretive power by providing the opportunity to characterize coupling between the metabolome and functional traits that held true not only within, but also among, tropical and temperate plant species.

### Plant multidimensional space

Following^7^, we explored constraints on the multidimensional space occupied by metabolic and classical functional traits separately for tropical and temperate species using an observed *versus* simulated hypervolume approach. Briefly, we calculated an *n*-dimensional convex hull volume (package “geometry”^100^) representing the observed multidimensional trait space occupied by species, where *n* is the number of metabolic (n = 5) or classical (n = 8) functional traits. We compared the observed hypervolume to the same four simulated null hypervolumes used by ref. ^7^, which tested the following null hypotheses: (i) traits vary independently and have uniform distributions (i.e., each trait is a unique axis and no selection against extreme trait values); (ii) traits vary independently but have normal distributions (i.e., each trait is a unique axis and selection against extreme trait values); (iii) traits vary independently but distributions are as observed (i.e., each trait is a unique axis but no assumptions about selection on values); and (iv) traits covary as but have normal distributions (i.e., no assumptions about trait interdependence but selection against extreme values). We simulated null hypervolumes 999 times and used permutation tests (package “ade4”^100^) to test for differences between the overall size of observed *versus* simulated hypervolumes, and presented percent changes between observed and mean simulated hypervolume sizes (where positive = observed > simulated; Table S2). We also calculated the “lumpiness” (i.e., aggregation around certain values) of species in multidimensional space by splitting *n*-dimensional space into 10 bins (i.e., 10^*n*^ cells), counting the number of species present in each cell, and calculating the minimum number of cells required to capture 10% of species^7^. Again, we tested for differences between observed and simulated null hypervolumes using permutation tests and presented percent changes between observed and mean simulated hypervolumes (Table S3).

### Axes of leaf metabolic specialization

We characterized major axes of leaf metabolic specialization separately among tropical or temperate species using varimax-rotated PCAs performed on metabolic functional traits. The first two PCs explained 92% variation for both tropical and temperate species, respectively, which we displayed using biplots of PC scores (Fig. 3). We explored whether axes of leaf metabolic specialization (Fig. 3) were orthogonal to, or colinear with, axes of classical functional trait variation (Fig. 4) by doing varimax-rotated PCAs containing metabolic plus classical functional traits, again separately among tropical and temperate species. Four PCs were needed to explain 66% and 67% variation among tropical and temperate species, respectively, which we displayed in two ways. First, we created a biplot matrix of all pairwise combinations of the first four axes (Fig. S3) to visualize the lack of overlap between metabolic and classical functional traits. Second, we performed Pearson correlations between PC scores and individual functional traits (Fig. 4, lower panels). Finally, we examined interdependence within and among metabolic and classical functional traits by performing pairwise Pearson correlations (Fig. 4, upper panels), clustering correlation coefficients (hierarchical clustering; Ward), and identifying discrete clusters of traits using the maximal silhouette width^96^ (Fig. 4, dendrograms). Dendrograms are organized based on the outcome of a tanglegram that tested for correspondence between the clustering of tropical and temperate traits (Fig. 4, central connectors).

### Data and materials availability

Raw tropical metabolomics data are available under the DOI 10.25345/C59J97^36^. Raw temperate species metabolomics data are available under the DOI 10.6084/m9.figshare.13032740^37^. Flat data and R code is available under the DOI 10.17605/OSF.IO/YZR4C.

## Acknowledgements

Special thanks to Jeanne Clément and Ghislain Vieilledent for building bespoke functionality into the R package “jSDM” for exploratory analyses. This study was part funded by a grant from the Synthesis Centre (sDiv) of the German Centre for Integrative Biodiversity Research (iDiv), awarded to TWNW, FS and NMvD. Plant material collection and metabolomics analysis were financed by Swiss National Science Foundation grants awarded to LP (grant no. 31003A-162604), SR (grant no. 31003A-131956), and JLW (grant no. CRSII5-189921/1). JMA is funded by the Swiss National Science Foundation (grant no. 31003A-176044). FS acknowledges funding by the University of Nottingham Anne McLaren Fellowship. ED and SR are funded by the Swiss National Science Foundation (grant no. 31003A_179481). VEJJ is funded by the French National Research Agency (grant no. ANR-17-CE01-0007: Mixxopeat). VB is supported by the French Agence Nationale pour la Recherche (grant no. ANR-12-BSV7-0016-01: SecPriMe2), the BioDivMeX Mistrals program, the Aix-Marseille University ‘Investissements d’Avenir’ program and the Labex OT-Med programs. PC is supported by a Ramón Areces Foundation Postdoctoral Scholarship. PDC is funded by the US National Science Foundation (grant no. DEB-1135733). NMvD, SM and KP are supported by iDiv Halle-Jena-Leipzig, which is funded by the German Research Foundation (grant nos DFG–FZT 118, 202548816). SN and KP are additionally funded by the German Network for Bioinformatics Infrastructure (de.NBI) and acknowledge BMBF funding (grant no. 031L0107). DLF is supported by the US National Science Foundation Graduate Research Fellowships Program. RSG is funded by a UK Natural Environment Research Council Independent Research Fellowship (grant no. NE/M018458/1). JP and JS are funded by the Spanish Government (grant no. CGL2016-79835), Catalan Government (grant no. SGR 2017-1005) and a European Research Council Synergy Grant (grant no. ERC-SyG-2013-610028: IMBALANCE-P). MCS is supported by a NOMIS Foundation project (“Remotely Sensing Ecological Genomics”) and by the University of Zürich Research Priority Program on Global Change and Biodiversity.

## Author contributions

Conceptualization: TWNW in collaboration with PMA, RDB, PC, BD, ED, AG-G, VEJJ, JP, SR, RSG, JS, MCS, FS, NMvD and WW, with support from all co-authors. Funding acquisition: FS, NMvD, and TWNW. Data collection: PMA, ED, and BD. Data curation: TWNW, with support from PMA, OB, PC, and ED. Formal analysis: TWNW, with support from PC, ED, MJE, DF, and VEJJ. Writing: TWNW, with support from MCS. Reviewing and editing: all co-authors.

## Materials & correspondence

Requests for materials and correspondence should be sent to the corresponding author.

## Extended Data Figures

**Extended Data Figure 1.**
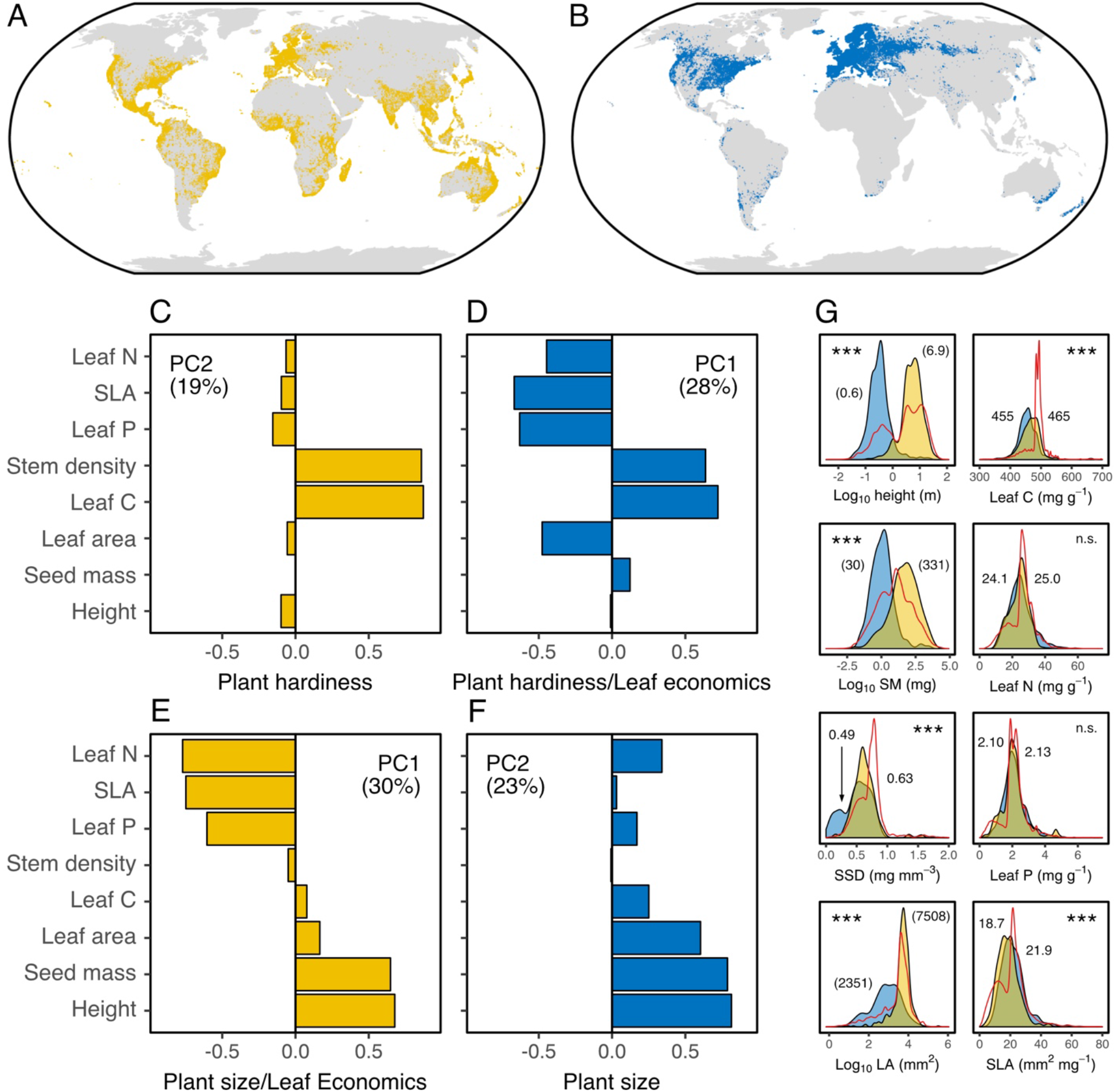
Tropical and temperate species capture major trade-offs in plant functional traits. (A,B) The global geographic extents of (A) tropical (yellow; N = 457) and (B) temperate (blue; N = 405) plant species used in this study. Points are processed GBIF records representing native and non-native occurrences, with sample collection being restricted to *in situ* native populations. (C-F) Loadings for the first two PCs of PCAs performed on eight classical functional traits separately for (C,E) tropical and (D,F) temperate species. Axes collectively capture well-established life-history trade-offs (i.e., leaf economics spectrum^6^, plant size/longevity^7,101^, plant hardiness^15^; see Main Text), although the coupling of these trade-offs and the ordering of PC axes differs between tropical and temperate species. (G) Kernel density estimates showing the distributions of classical traits used in PCAs (height, seed mass (SM), stem specific density (SSD), leaf area (LA) and leaf carbon, nitrogen and phosphorus) for tropical (blue) and temperate (yellow) species, as well as for all species in the TRY database (red ribbon^89^). Text shows group means, with values in brackets being back-transformed to original units (log_10_-transformed traits; see Methods). Asterisks illustrate significant differences *(P* < 0.001; linear models) between tropical and temperate species means.

