## Supplementary Material for "Leaf metabolic traits reveal hidden dimensions of plant form and function"

--- SUPPLEMENTARY INFORMATION ---

Tom W. N. Walker, Franziska Schrodt, Pierre-Marie Allard, Emmanuel Defossez, Vincent E. J. Jassey, Mereditch C. Schuman, Jake M. Alexander, Oliver Baines, Virginie Baldy, Richard D. Bardgett, Pol Capdevila, Phyllis D. Coley, Nicole M. van Dam, Bruno David, Patrice Descombes, Maria-Jose Endara, Catherine Fernandez, Dale Forrister, Albert Gargallo-Garriga, Gaëtan Gauser, Sue Marr, Steffen Neumann, Loïc Pellissier, Kristian Peters, Sergio Rasmann, Ute Roessner, Roberto Salguero-Gómez, Jordi Sardans, Wolfram Weckwerth, Jean-Luc Wolfender & Josep Peñuelas

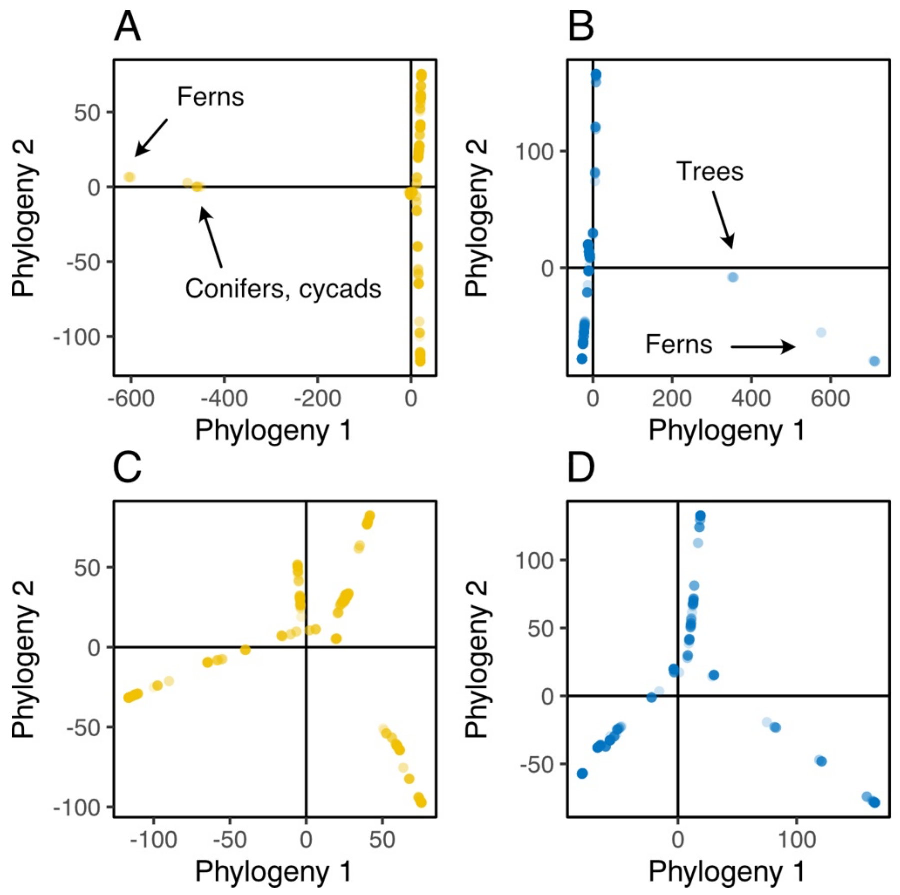


**Supplementary Figure S1 I Ordinations of phylogenetic distances.** Biplots of scores from the first two axes of principal coordinates analyses performed on cophenetic phylogenetic distances of (A,C) tropical (yellow) and (B,D) temperate (blue) species, either (A,B) with (tropical N = 471, temperate N = 414) or (C,D) without (tropical N = 457, temperate N = 405) phylogenetically distinct species present. Subsetted data (C,D) were used for downstream analyses.

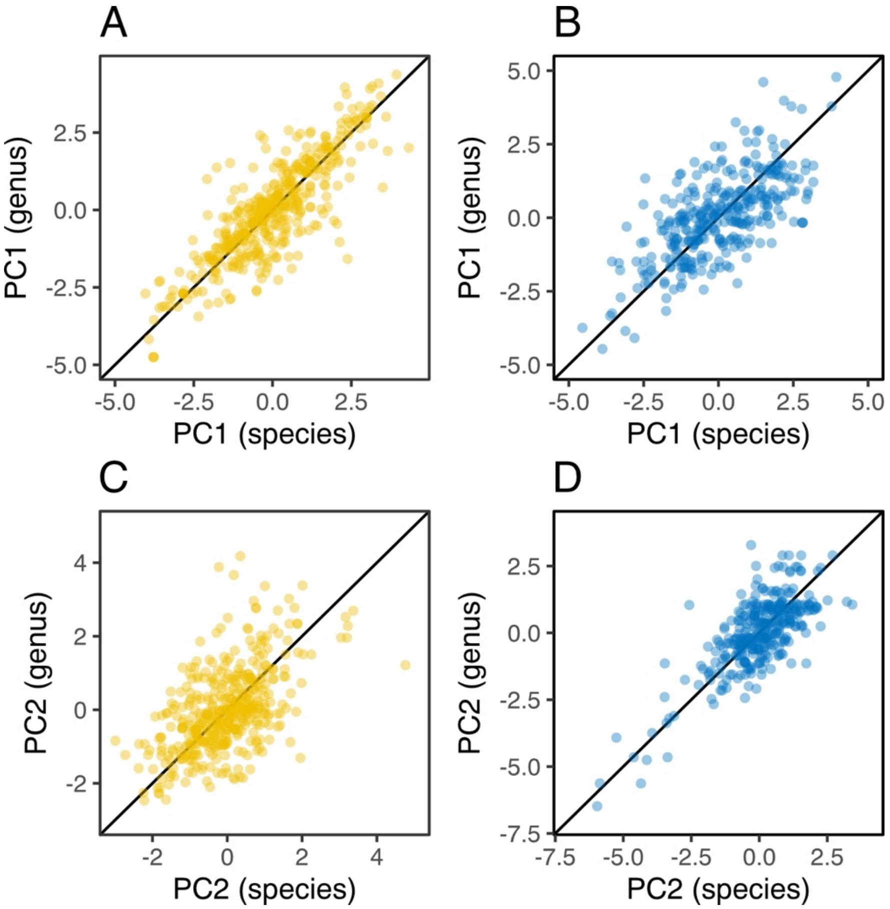


**Supplementary Figure S2 | Using genus-level functional traits does not change nature of resulting PC axes.** Correlations between (A,B) PC1 and (C,D) PC2 scores from PCAs performed on classical functional traits calculated at the species versus (x-axis) genus (y-axis) level separately for tropical (yellow) and temperate (blue) species. A 1:1 line is displayed for reference.

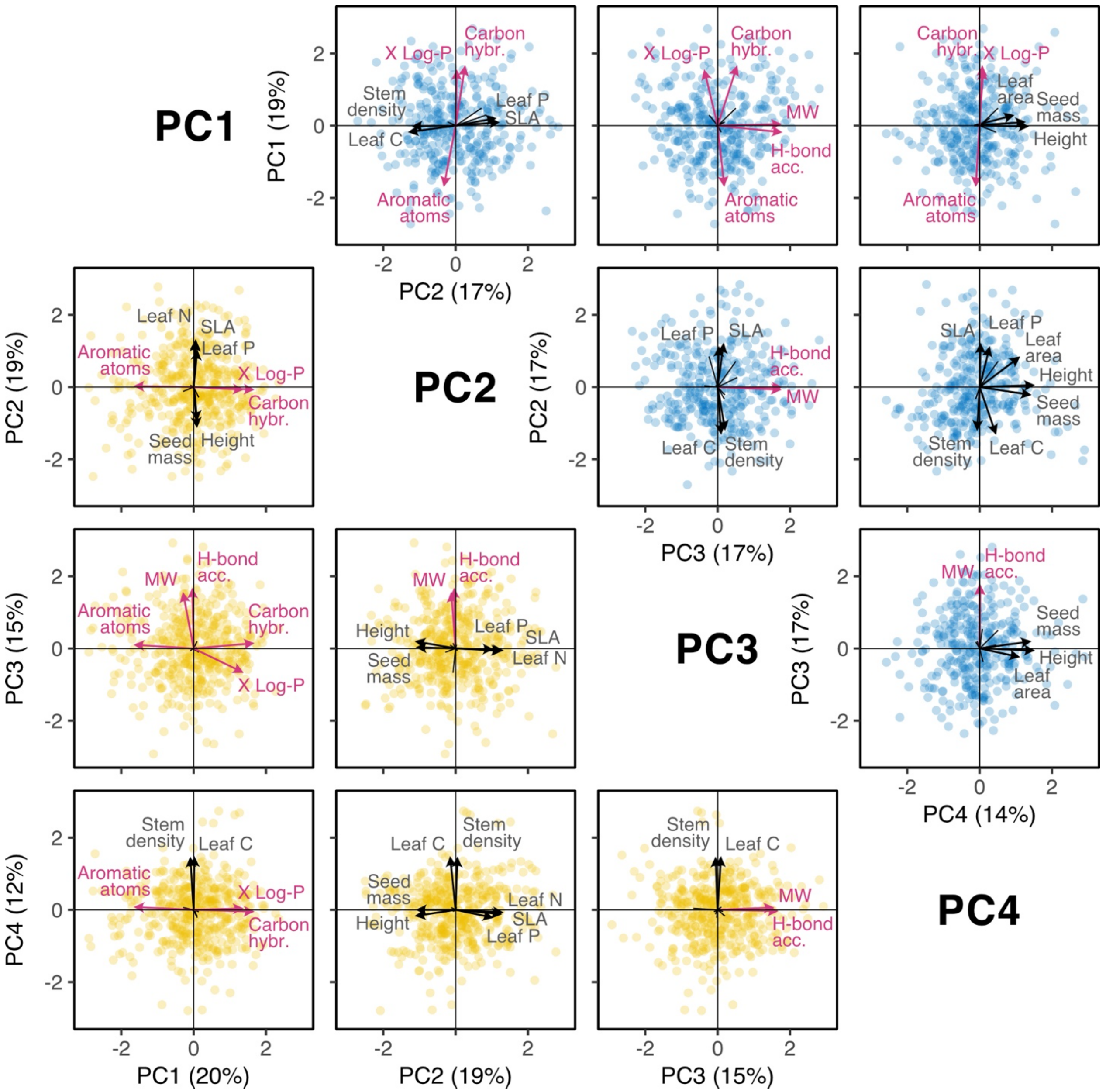


**Supplementary Figure S3 | Metabolomic and classical functional traits are always orthogonal.** A scatterplot matrix showing biplots of all combinations of the first four axes of PCAs performed on metabolomic (red) plus classical (grey) functional traits for tropical (bottom left, yellow; N = 457) or temperate (top right, blue; N = 405) species. Points show positions of species on the two axes, while arrows show the strength and direction of trait loadings.

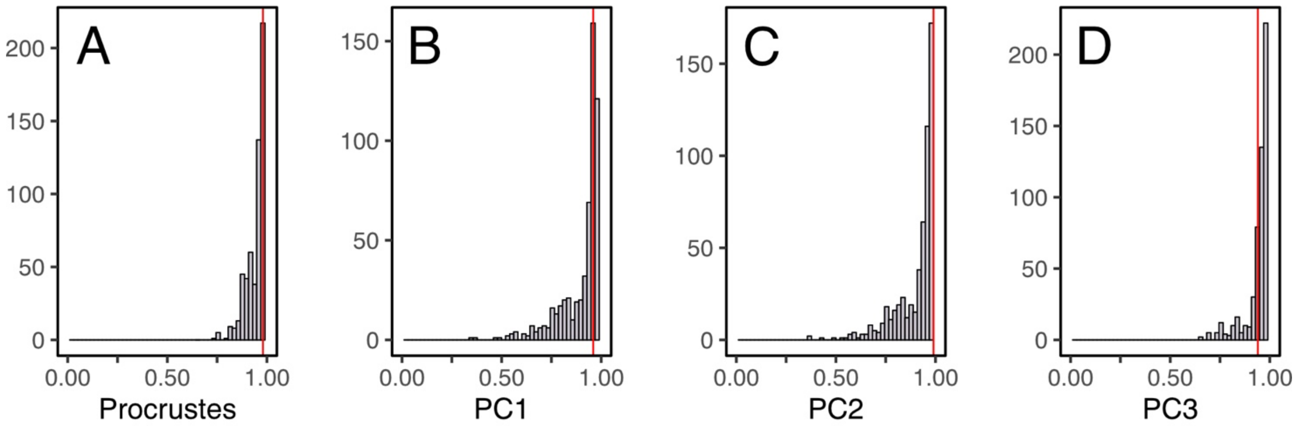


**Supplementary Figure S4 | Subset PCAs are equivalent to the full PCA for most combinations of properties.** Histograms of correlation coefficients between a full PCA containing all 21 chemical properties and subset PCAs performed on every combination of five chemical properties that cover the five clusters of leaf chemical variation (i.e. one from each cluster; N = 576). (A) Procrustes Rotation tests among contributing distance matrices. (B-D) Pairwise Pearson correlations among PC scores, where PCs from subset PCAs are first matched to corresponding PCs from the full PCA (i.e., allowing for the ordering of PCs to change among subset PCAs; *x*-axis). Red lines illustrate correlation coefficients for the subset PCA with selected chemical properties (see Text).

**Table S1 l Selected chemical properties.** Names, descriptions and units of 21 quantitative chemical properties used to characterise leaf metabolite chemistry, as well as their observed ranges across all unique annotated metabolites from tropical and temperate species (see Main Text). Chemical properties were derived from the CDK [1] using SMILES chemical identifiers (Methods).

| Category | Property | Description | Unit | Data range |
|---|---|---|---|---|
| Constitutional | Molecular weight (MW) | Total metabolite mass, as calculated from masses of constituent atoms | Da | 80.1 – 3158.7 |
| Constitutional | Total atom count | Total number of atoms | # | 5 – 425 |
| Constitutional | Aromatic atom count | Number of atoms in aromatic rings; high in shikimate pathway derivatives (e.g., flavonoids) [2] | # | 0 – 36 |
| Constitutional | Longest chain atom count | Number of atoms in longest chain; high in lipids [3] | # | 3 – 53 |
| Constitutional | Largest pi-system atom count | Number of atoms in largest conjugated system (i.e., alternating double-single bonds); high in pigments [4] and light-absorbing compounds [5] | # | 0 – 42 |
| Constitutional | Total bond count | Total number of (non-H) bonds | # | 2 – 223 |
| Constitutional | Aromatic bond count | Number of (non-H) bonds in aromatic rings; high in shikimate pathway derivatives (e.g., flavonoids) [2] | # | 0 – 40 |
| Constitutional | Rotatable bond count | Number of rotatable bonds (i.e., single, non-ring, non-terminal bonds with low energy barrier for rotation); negatively correlates with passive transport across biological membranes [6] | # | 0 – 86 |
| Constitutional | X log *P* | Octanol/water partition coefficient calculated using a modified atom-additive model summing atomic contributions and correcting for intramolecular interactions [7]; positive indicates affinity to octanol (i.e., nonpolar, hydrophobic), negative to water (i.e., polar, hydrophilic) | Log-ratio | -12.7 – 32.1 |
| Constitutional | A log *P* | Octanol/water partition coefficient calculated using an atom-additive model summing atom type contributions based on focal atom and bond characteristics [8]; positive indicates affinity to octanol (i.e., nonpolar, hydrophobic), negative to water (i.e., polar, hydrophilic) | Log-ratio | -15.1 – 27.2 |
| Constitutional | M log *P* | Octanol/water partition coefficient calculated using a simple equation dependent on the number of C atoms and number of hetero atoms [9]; smaller indicates affinity to octanol (i.e., nonpolar, hydrophobic), larger to water (i.e., polar, hydrophilic) | Log-ratio | 0.7 – 8.3 |
| Topological | Topological polar surface area (PSA) | Sum of surface area of polar atoms, calculated from contributions of polar molecular fragments [10]; in pharmaceutical studies, PSA correlates inversely with passive transport through biological membranes (lower TPSA means greater fraction absorbed) [11] | Å | 0 – 1468.2 |
| Topological | MW-specific PSA | PSA divided by metabolite molecular mass; in pharmaceutical studies, PSA correlates inversely with passive transport through biological membranes [11] | Å $Da^{-1}$ | 0 – 0.77 |
| Topological | Hybridisation ratio | Fraction of $sp_3$ to $sp_2$ carbon atoms; proxy for bond saturation and three-dimensional topological complexity; positive correlate of melting point, solubility, and bioactivity [12] | Ratio | 0 – 1 |
| Topological | Fractional $CSP_3$ | Fraction of $sp_3$ carbon atoms to total carbon count; proxy for bond saturation and three-dimensional topological complexity; positive correlate of melting point, solubility, and bioactivity [12] | Ratio | 0 – 1 |
| Topological | $f_{MF}$ | Ratio between size of molecular framework (ring atoms plus linkers) and size of metabolite; complexity measure positively correlated to promiscuity (number of protein targets ≥50% inhibited) at values above 0.65 [13] | Ratio | 0 – 1 |
| Topological | Eccentric connectivity index | Distance-cum-adjacency topological descriptor (higher for longer chains with less branching); correlates with size and physicochemical properties (e.g., boiling point) [14] | - | 6 – 14213 |
| Topological | Wiener path number | Topological descriptor of molecular branching that can differentiate structural isomers; correlates positively with size and boiling point [15] | # | 4 – 413341 |
| Topological | Wiener polarity number | A variant of Wiener path number calculated using vertices (C atoms) at distance 3; correlates positively with size and boiling point [15] | # | 0 – 354 |
| Electronic | H-bond donor count | Number of H-bond donors (OH/NH, formal charge ≥ 0); H-bonds are intermolecular forces essential for macromolecules and complexes [16] | # | 0 – 44 |
| Electronic | H-bond acceptor count | Number of H-bond acceptors (O/N, formal charge ≤ 0, non-ether O, non-adjacent ON); H-bonds are intermolecular forces essential for macromolecules and complexes [16] | # | 0 – 75 |

**Table S2 | Relative differences between the sizes of hypervolumes for four null models and those observed for metabolic or classical functional traits.** Percent differences between mean null model hypervolume sizes (999 permutations) versus observed hypervolumes for metabolic (MT) or classical (FT) functional traits. Values not in parentheses are statistically significant (underlined: $P < 0.05$; others: $P < 0.01$).

| **Null model** | | **Tropical species** | | **Temperate species** | |
|---|---|---|---|---|---|
| Trait distribution | Trait covariance | MTs | FTs | MTs | FTs |
| Uniform | Independent | - 99.8% | - 99.9% | - 99.8% | - 99.5% |
| Normal | Independent | - 98.2% | - 52.7% | - 98.1% | - 56.6% |
| As observed | Independent | - 98.3% | - 48.9% | - 98.1% | - 53.7% |
| Normal | As observed | + 27.7% | (- 6.5%) | + 15.3% | - 15.2% |

**Table S3 | Relative differences between the lumpiness of hypervolumes for four null models and those observed for metabolic or classical functional traits.** Mean percent differences between the minimum number of cells in multidimensional space needed to cover 10% species for four null model hypervolumes (999 permutations) versus observed hypervolumes for metabolic (MTs) or classical (FT) functional traits. Values not in parentheses are statistically significant ($P < 0.01$).

| **Null model** | | **Tropical species** | | **Temperate species** | |
|---|---|---|---|---|---|
| Trait distribution | Trait covariance | MTs | FTs | MTs | FTs |
| Uniform | Independent | - 94.7% | - 44.1% | - 89.2% | - 26.4% |
| Normal | Independent | - 83.6% | - 65.2% | - 71.4% | - 22.6% |
| As observed | Independent | - 81.6% | - 57.2% | - 71.0% | - 19.2% |
| Normal | As observed | (- 15.8%) | - 62.7% | (+ 5.2%) | - 18.9% |

**Table S4 | Classical functional trait coverage.** Percent coverage for the eight classical functional traits used here in the original TRY dataset and following 90 iterations of BHPMF imputations. In both cases, clear outliers were removed (see Methods).

| **Trait** | **Tropical species** | | **Temperate species** | |
|---|---|---|---|---|
| | **No imputation** | **BHPMF** | **No imputation** | **BHPMF** |
| Plant height | 41.6% | 81.6% | 99.7% | 99.7% |
| Seed mass | 67.2% | 87.1% | 92.0% | 92.3% |
| Stem density | 50.3% | 58.2% | 17.7% | 27.4% |
| Leaf area | 45.7% | 69.1% | 88.2% | 89.7% |
| Specific leaf area | 48.1% | 57.1% | 91.2% | 94.4% |
| Leaf carbon content | 32.8% | 46.8% | 75.8% | 80.2% |
| Leaf nitrogen content | 45.3% | 63.2% | 79.9% | 91.4% |
| Leaf phosphorus content | 40.7% | 62.6% | 56.0% | 94.1% |
